# SARS-CoV-2 Infection Depends on Cellular Heparan Sulfate and ACE2

**DOI:** 10.1101/2020.07.14.201616

**Authors:** Thomas Mandel Clausen, Daniel R. Sandoval, Charlotte B. Spliid, Jessica Pihl, Chelsea D. Painter, Bryan E. Thacker, Charles A. Glass, Anoop Narayanan, Sydney A. Majowicz, Yang Zhang, Jonathan L. Torres, Gregory J. Golden, Ryan Porell, Aaron F. Garretson, Logan Laubach, Jared Feldman, Xin Yin, Yuan Pu, Blake Hauser, Timothy M. Caradonna, Benjamin P. Kellman, Cameron Martino, Philip L.S.M. Gordts, Sandra L. Leibel, Summit K. Chanda, Aaron G. Schmidt, Kamil Godula, Joyce Jose, Kevin D. Corbett, Andrew B. Ward, Aaron F. Carlin, Jeffrey D. Esko

## Abstract

We show that SARS-CoV-2 spike protein interacts with cell surface heparan sulfate and angiotensin converting enzyme 2 (ACE2) through its Receptor Binding Domain. Docking studies suggest a putative heparin/heparan sulfate-binding site adjacent to the domain that binds to ACE2. In vitro, binding of ACE2 and heparin to spike protein ectodomains occurs independently and a ternary complex can be generated using heparin as a template. Contrary to studies with purified components, spike protein binding to heparan sulfate and ACE2 on cells occurs codependently. Unfractionated heparin, non-anticoagulant heparin, treatment with heparin lyases, and purified lung heparan sulfate potently block spike protein binding and infection by spike protein-pseudotyped virus and SARS-CoV-2 virus. These findings support a model for SARS-CoV-2 infection in which viral attachment and infection involves formation of a complex between heparan sulfate and ACE2. Manipulation of heparan sulfate or inhibition of viral adhesion by exogenous heparin may represent new therapeutic opportunities.

## Introduction

The COVID-19 pandemic, caused by the novel respiratory coronavirus 2 (SARS-CoV-2), has swept across the world, resulting in serious clinical morbidities and mortality, as well as widespread disruption to all aspects of society. As of July 10^th^, 2020, the virus has spread to 213 countries, causing more than 12.3 million confirmed infections and at least 555,000 deaths (World Health Organization). Current isolation/social distancing strategies seek to flatten the infection curve to avoid overwhelming hospitals and to give the medical establishment and pharmaceutical companies time to develop and test antiviral drugs and vaccines. Currently, only one antiviral agent, Remdesivir, has been approved for adult COVID-19 patients (Beigel et al., 2020) and vaccines may be 12-18 months away. Understanding the mechanism for SARS-CoV-2 infection and its tissue tropism could reveal other targets to interfere with viral infection and spread.

The glycocalyx is a complex mixture of glycans and glycoconjugates and is continuous with the extracellular matrix. Given its location, viruses and other infectious organisms, must pass through the glycocalyx to engage receptors thought to mediate viral entry into host cells. Many viral pathogens utilize glycans as attachment factors to facilitate the initial interaction with host cells, including influenza virus, Herpes simplex virus, human immunodeficiency virus, and different coronaviruses (SARS-CoV-1 and MERS) (Cagno et al., 2019; Koehler et al., 2020; Stencel-Baerenwald et al., 2014). Several viruses interact with sialic acids, which are located on the ends of glycans found in glycolipids and glycoproteins (Varki et al., 2015). Other viruses interact with heparan sulfate (HS) (Milewska et al., 2014), a highly negatively charged linear polysaccharide that is attached to a small set of membrane or extracellular matrix proteoglycans (Lindahl et al., 2015). In general, glycan-binding domains on membrane proteins of the virion envelope mediate initial attachment of virions to glycan receptors. Attachment in this way can lead to the engagement of protein receptors on the host plasma membrane that facilitate membrane fusion or engulfment and internalization of the virion.

Like other macromolecules, HS can be divided into subunits, which are operationally defined as disaccharides based on the ability of bacterial enzymes or nitrous acid to cleave the chain into disaccharide units (Esko and Selleck, 2002). The basic disaccharide subunit consists of □1-4 linked D-glucuronic acid (GlcA) and □1-4 linked *N*-acetyl-d-glucosamine (GlcNAc), which undergo various modifications by sulfation and epimerization while the copolymer assembles on a limited number of membrane and extracellular matrix proteins (only 17 heparan sulfate proteoglycans are known) (Lindahl et al., 2015). The variable length of the modified domains and their pattern of sulfation create unique motifs to which HS-binding proteins interact (Xu and Esko, 2014). Different tissues and cell types vary in the structure of HS, and HS structure can vary between individuals and with age (de Agostini et al., 2008; Feyzi et al., 1998; Han et al., 2020; Ledin et al., 2004; Vongchan et al., 2005; Warda et al., 2006; Wei et al., 2011). These differences in HS composition may contribute to the tissue tropism of viruses and other pathogens.

In this report, we show that the ectodomain of the SARS-CoV-2 spike (S) protein interacts with cell surface HS through the Receptor Binding Domain (RBD) in the S1 subunit. Binding of SARS-CoV-2 S protein to cells requires engagement of both cellular HS and angiotensin converting enzyme 2 (ACE2), suggesting that HS acts as a coreceptor. Therapeutic unfractionated heparin (UFH), non-anticoagulant heparin and HS derived from human lung and other tissues blocks binding. UFH and heparin lyases also block infection of cells by S protein pseudotyped virus and native SARS-CoV-2. These findings identify cellular HS as a necessary co-factor for SARS-CoV-2 infection and emphasizes the potential for targeting S protein-HS interactions to attenuate virus infection.

## Results

### Molecular modeling reveals an HS-binding site adjacent to the ACE2 binding domain in the SARS-CoV-2 spike protein

The trimeric SARS-CoV-2 S protein is thought to engage human ACE2 with a single RBD extending from the trimer in an “open” active conformation (Fig. 1A) (Walls et al., 2020; Wrapp et al., 2020). Adjacent to the ACE2 binding site and exposed in the open RBD conformation lies a group of positively-charged amino acid residues that represents a potential site that could interact with heparin or heparan sulfate (Fig. 1A and Suppl. Fig. S1). We calculated an electrostatic potential map of the RBD (from PDB ID 6M17 (Yan et al., 2020)), which revealed an extended electropositive surface with dimensions and turns/loops consistent with a heparin-binding site (Fig. 1B) (Xu and Esko, 2014). Docking studies using a tetrasaccharide (dp4) fragment derived from heparin demonstrated preferred interactions with this electropositive surface, which based on its dimensions could accommodate a chain of up to 20 monosaccharides (Fig. 1B and 1C). Evaluation of heparin-protein contacts and energy contributions using the Molecular Operating Environment (MOE) software suggested strong interactions with the positively charged amino acids R346, R355, K444, R466 and possibly R509 (Figs. 1A, 1D, and 1E). Other amino acids, notably F347, S349, N354, G447, Y449, and Y451, could coordinate the oligosaccharide through hydrogen bonds and hydrophobic interactions. Notably, the putative binding surface for oligosaccharides is adjacent to, but separate from the ACE2 binding site, suggesting that a single RBD could simultaneously bind both cell-surface HS oligosaccharides and its ACE2 protein receptor. Consistent with the hypothesis that only one RBD in the trimeric S protein is exposed in an active conformation, the putative HS binding is partially obstructed in the closed RBD conformation, while exposed in the active state (Suppl. Fig. S1).

**Figure 1.**
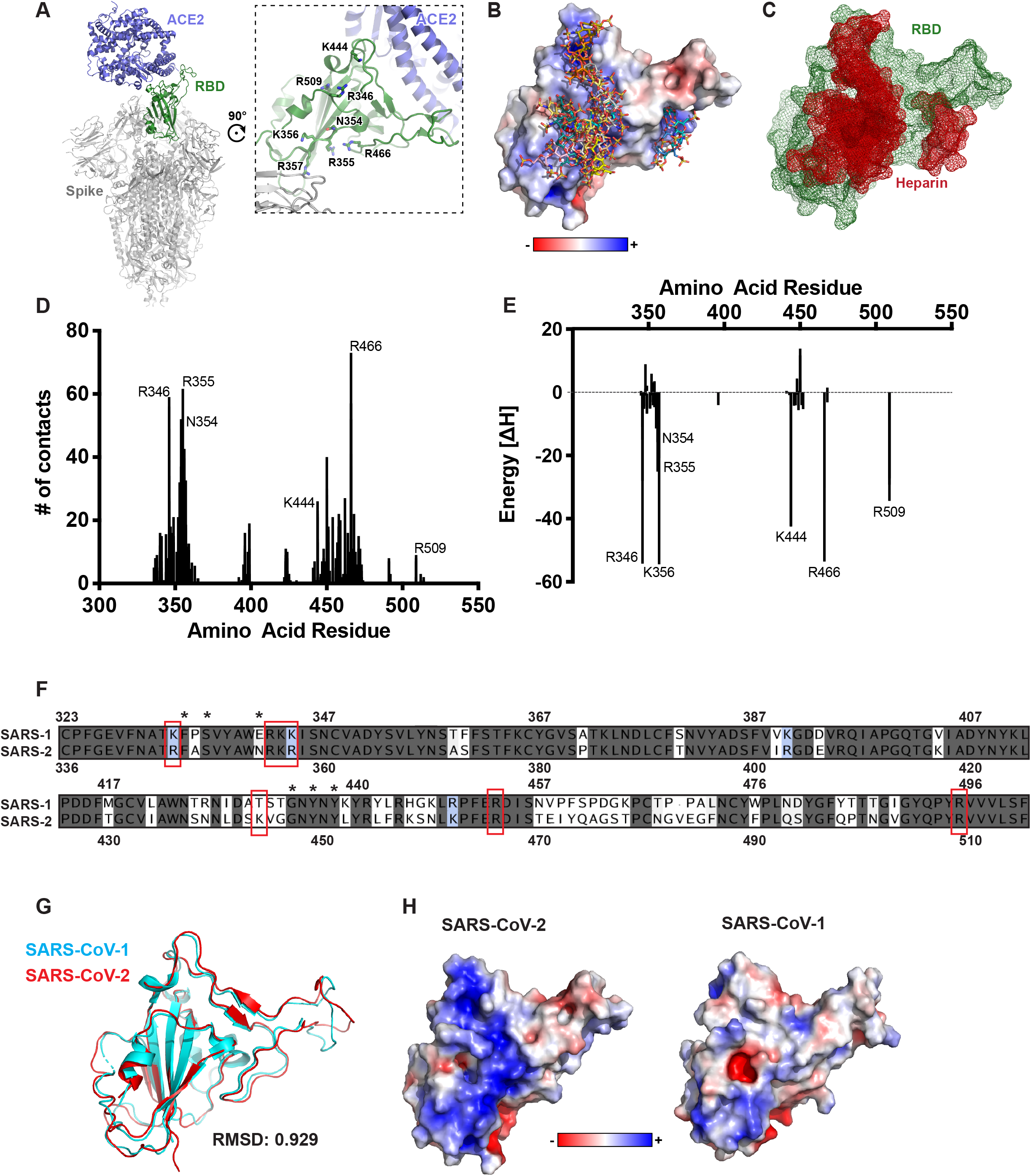
Molecular modeling of the SARS-Cov-2 spike RBD interaction with heparin. **A**, A molecular model of SARS CoV-2 S protein trimer (PDB: 6VSB and 6M0J) rendered with Pymol. ACE2 is shown in blue and the RBD in open conformation in green. A set of positively-charged residues lies distal to the ACE2 binding site. **B**, Electrostatic surface rendering of the SARS-CoV-2 RBD (PDB: 6M17) docked with dp4 heparin oligosaccharides. Blue and red surfaces indicate electropositive and electronegative surfaces, respectively. Oligosaccharides are represented using standard CPK format. **C,** Mesh surface rendering of the RBD (green) docked with dp4 heparin oligosaccharides (red). **D**, Number of contacts between the RBD amino acids and a set of docked heparin dp4 oligosaccharides from **A** and **B. E,** Calculated energy contributions of each amino acid residue in the RBD that can interact with heparin. **F,** Amino acid sequence alignment of the SARS-CoV-1 and SARS-Cov-2 RBD. Red boxes indicate amino acid residues contributing to the electropositive patch in **A and C.** Identical residues are shaded blue. Conservative substitutions have backgrounds in shades of pink. Non-conserved residues have a white background **G,** Structural alignment of SARS-CoV-1 (cyan; PDB:3GBF) and SARS-CoV-2 RBDs (red; PDB: 6M17) RBD. **H,** Electrostatic surface rendering of the SARS-CoV-1 and SAR-CoV-2 RBDs.

The S protein RBD of SARS-CoV-2 S is 73% identical with the same domain of SARS-CoV-1 S (Fig. 1F). These domains are highly similar in structure with an overall Cα r.m.s.d. of 0.929 Å (Fig. 1G). However, an electrostatic potential map of the SARS-CoV-1 S RBD does not show an electropositive surface like that observed in SARS-CoV-2 (Fig. 1H). Most of the positively charged residues comprising this surface are conserved between the two proteins, with the exception of SARS-CoV-2 K444 which is a threonine in SARS-CoV-1 (Fig. 1F and G). Additionally, the other amino acid residues predicted to coordinate with the oligosaccharide are conserved with the exception of Asn354 in SARS-CoV-2, which is a glutamate residue in SARS-CoV-1. SARS-CoV-1 has been shown to interact with cellular HS in addition to its entry receptors ACE2 and transmembrane protease, serine 2 (TMPRSS2) (Lang et al., 2011). Our analysis suggests that the putative heparin-binding site in SARS-CoV-2 S may mediate an enhanced interaction with heparin compared to SARS-CoV-1, and that this change evolved through as few as two amino acid substitutions, Thr→Lys444 and Glu→Asn354.

### The SARS-CoV-2 spike protein binds heparin through the RBD domain

To test experimentally if the SARS-CoV-2 S protein interacts with heparin, recombinant ectodomain and RBD proteins were prepared and characterized. Initial studies encountered difficulty in stabilizing the S ectodomain protein, a problem that was resolved by raising the concentration of NaCl to 0.3 M in HEPES buffer. Under these conditions, the protein could be stored at room temperature, 4 °C or at −80 °C for at least two weeks. SDS-PAGE showed that each protein was ~98% pure (Fig. Suppl. S2A). Transmission electron micrographs of the S ectodomains revealed trimeric structures (Suppl. Fig. S2B). The main component by SEC behaved as a glycosylated trimer with an apparent molecular mass of ~740 kDa (Suppl. Fig. S2C). A highly purified commercial preparation of RBD protein was used in some studies (SINO Biologics, Suppl. Fig. S2A) as well as recombinant RBD protein expressed in ExpiHEK cells (Suppl. Fig. S2D), both of which were judged >98% pure by SDS-PAGE and SEC (Suppl. Fig. S2E).

Recombinant S ectodomain and RBD proteins were applied to a column of heparin-Sepharose. Elution with a gradient of sodium chloride showed that the RBD eluted at ~0.3 M NaCl, with a shoulder that eluted with higher salt (Fig. 2A). Recombinant S ectodomain also bound to heparin-Sepharose, but it eluted across a broader concentration of NaCl. The elution profiles suggest that the preparations contained a population of molecules that bind to heparin, but that some heterogeneity in affinity for heparin occurs, which may reflect differences in glycosylation, oligomerization or binding heparin binding sites in the closed conformation.

**Figure 2.**
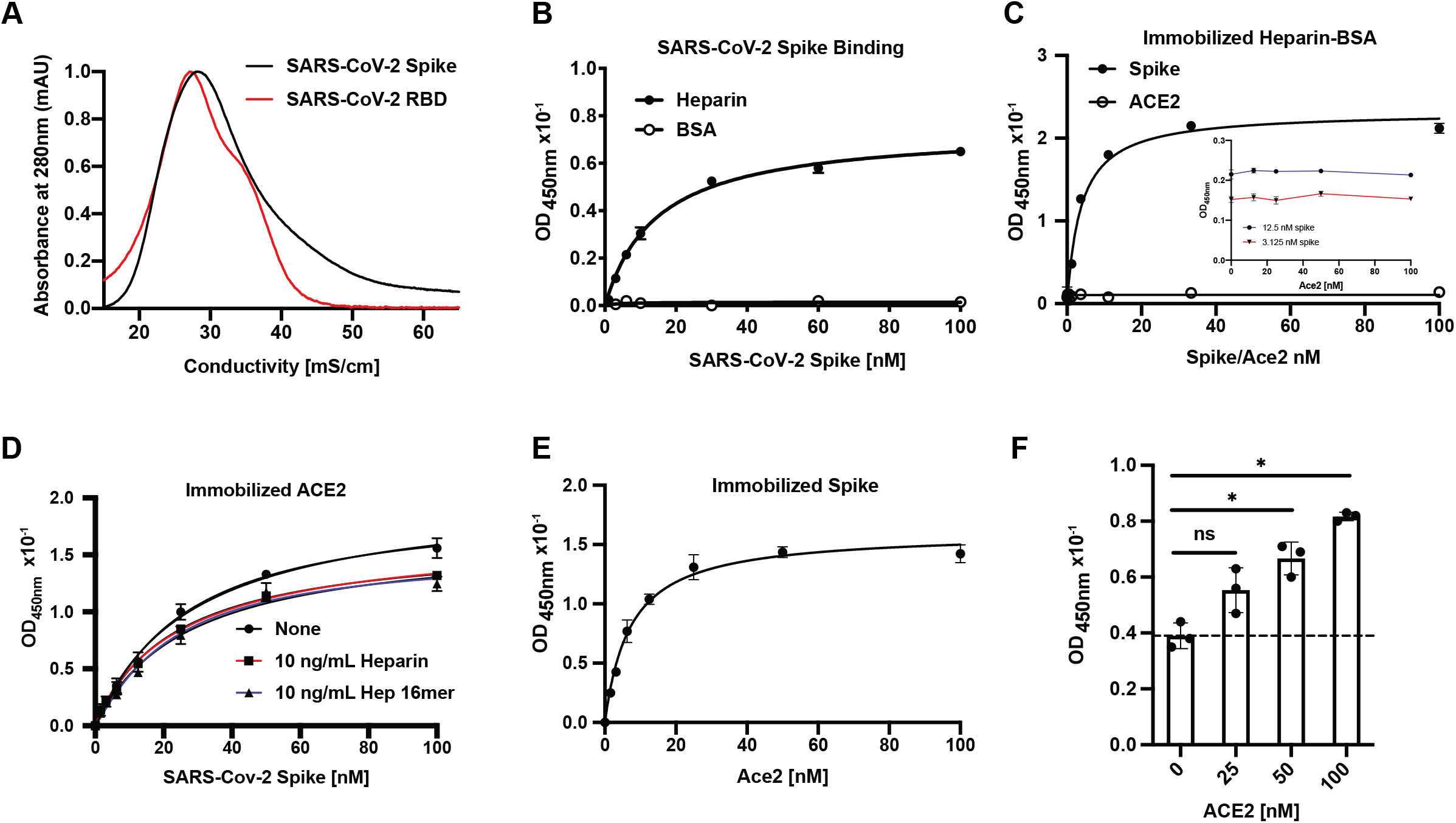
SARS-CoV-2 spike binds heparin through the RBD domain. **A**, Recombinant trimeric SARS-CoV-2 spike and RBD proteins were bound to heparin-Sepharose and eluted with a gradient of sodium chloride (broken line). **B,** Spike protein binds to immobilized unfractionated heparin. **C,** Binding of spike protein or ACE2 to heparin-BSA. Insert shows SARS-CoV-2 spike protein binding to heparin-BSA in the presence of ACE2. **D,** SARS-CoV-2 spike protein binding to immobilized recombinant ACE2 protein in the presence and absence of heparin or a heparin 16-mer. **E,** ACE2 binding to immobilized spike protein. **F,** ACE2 binding to spike protein immobilized on heparin-BSA. The broken line represents baseline binding. Statistical analysis was by one-way ANOVA. (ns: p > 0.05, *: p ≤ 0.05, **: p ≤ 0.01, ***: p ≤ 0.001, ****: p ≤ 0.0001).

The S ectodomain also bound in a concentration-dependent manner to unfractionated heparin immobilized on a plate, and the data fit a binding curve with an apparent K_*D*_ value of 15 ± 1 nM (Fig. 2B). S ectodomain protein also bound in a saturable manner to heparin-BSA conjugates immobilized on a plate (Fig. 2C). In contrast, ACE2 did not bind to heparin-Sepharose. ACE2 also had no effect on binding of S protein to heparin-BSA at all concentrations that were tested (Fig. 2C, inset). The ectodomain protein bound immobilized recombinant ACE2 (Fig. 2D). Inclusion of UFH or hexadecasaccharides (dp16) from heparin did not affect the affinity of S protein for ACE2 and only mildly decreased the extent of binding in this assay format (Fig. 2D). Biotinylated ACE2 bound to immobilized S protein (Fig. 2E). A ternary complex of heparin, ACE2 and S protein could be demonstrated by binding of S protein to immobilized heparin-BSA and titrating biotinylated ACE2. Binding of ACE2 under these conditions increased in proportion to the amount of S protein bound to the heparin-BSA (Fig. 2F). These findings support a model in which the binding sites for HS and ACE2 in the RBD act independently and can engage their receptors simultaneously.

### The SARS-CoV-2 spike protein depends on cellular heparan sulfate for cell binding

The assays described in Fig. 2 utilized heparin, which is often used as a surrogate for cellular HS due to its commercial availability. Although it is related in structure to HS, heparin is much more highly sulfated and negatively charged and can act as a strong unspecific cation exchanger. To determine if the SARS-CoV-2 S protein binds to typical HS found on cells, S ectodomain protein was added to human H1299 cells, an adenocarcinoma cell line derived from Type 2 alveolar cells (Fig. 3A). Spike ectodomains bound to H1299 cells, yielding an apparent K_D_ value of ~75 nM. Treatment of the cells with a mixture of heparin lyases (HSase), which degrades cell surface HS, dramatically reduced binding (Fig 3A). The S ectodomain also bound to human A549 cells, another Type 2 alveolar adenocarcinoma line, as well as human hepatoma Hep3B cells (Fig. 3B). Removal of HS by enzymatic treatment dramatically reduced binding in both of these cell lines as well (Fig. 3B). Recombinant RBD protein also bound to all three cell lines and binding largely depended on HS (Fig. 3C). A melanoma cell line, A375, was tested independently and also showed HS dependent binding (Fig 3D). The extent of binding to HS across the four cell lines varied ~4-fold. This variation was not due to differences in HS expression as illustrated by staining of cell surface HS with mAb 10E4, which recognizes a common epitope in HS (Fig. 3E). In fact, A375 cells have the highest expression of cell surface HS, but the lowest extent of S protein binding (Fig. 3D), whereas Hep3B cells have low amounts of cell surface HS and the highest binding of S protein. This lack of correlation could reflect differences in the structure of HS or available ACE2.

**Figure 3.**
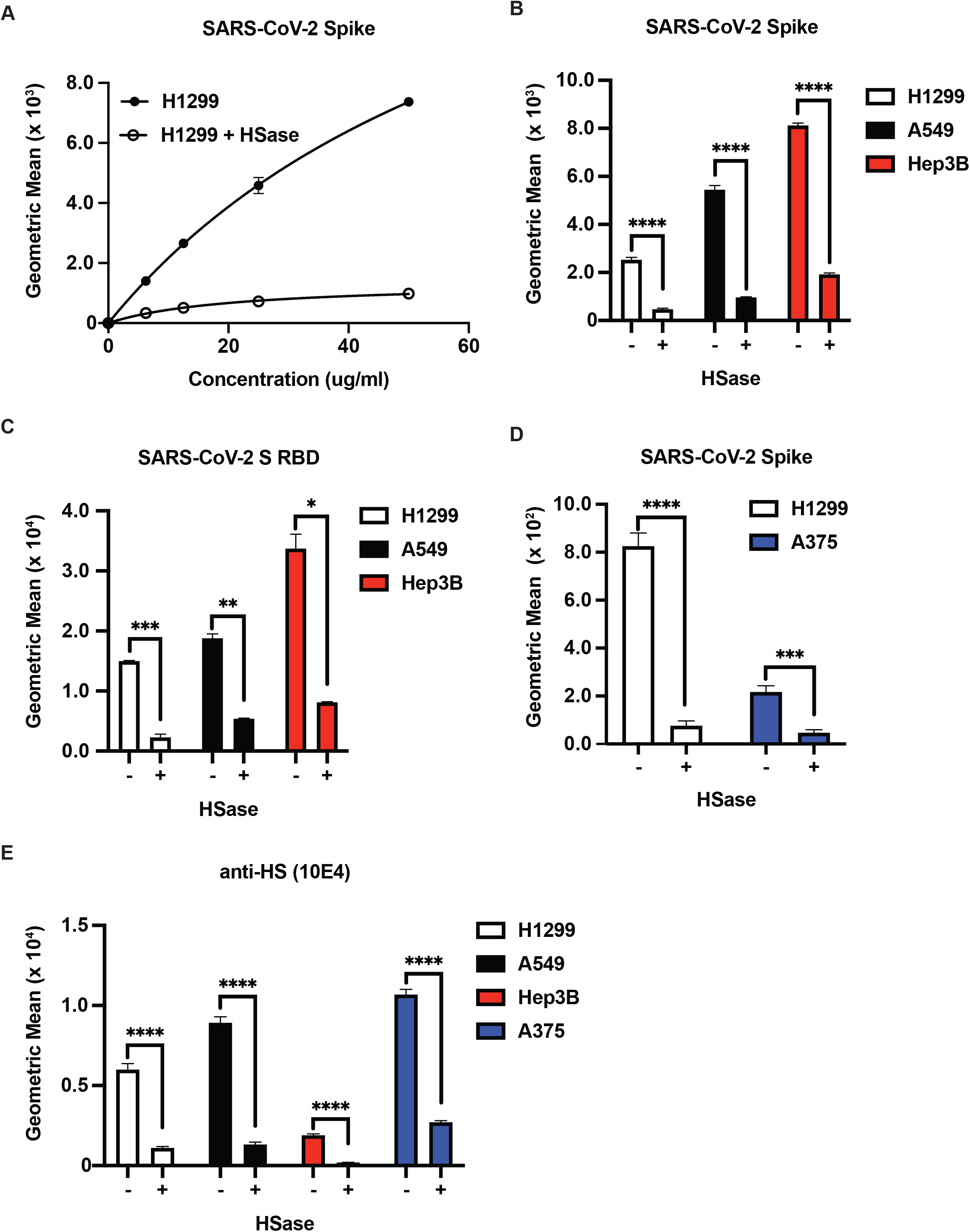
SARS-CoV-2 spike ectodomain binding to cells is dependent on cellular HS. **A,** Titration of recombinant SARS-CoV-2 spike protein binding to human H1299 cells with and without treatment with a mix of heparin lyases I, II, and III (HSase). **B,** Recombinant SARS-CoV-2 spike protein staining (20 μg/ml) of H1299, A549 and Hep3B cells, with and without HSase treatment. **C,** SARS-CoV-2 S RBD protein binding (20 μg/ml) to H1299, A549 and Hep3B cells with and without HSase treatment. **D,** SARS-CoV-2 spike protein binding (20 μg/ml) to H1299 and A375 cells with and without HSase treatment. **E,** Anti-HS (F58-10E4) staining of H1299, A549, Hep3B and A375 cells with and without HSase-treatment. All values were obtained by flow cytometry. Graphs shows representative experiments performed in technical triplicate. The experiments were repeated at least three times. Statistical analysis by unpaired t-test (ns: p > 0.05, *: p ≤ 0.05, **: p ≤ 0.01, ***: p ≤ 0.001, ****: p ≤ 0.0001).

We also measured binding of the S ectodomain and RBD proteins to a library of mutant Hep3B cells, carrying CRISPR/Cas9 induced mutations in biosynthetic enzymes essential for synthesizing HS (Anower et al., 2019). Inactivation of *EXT1*, a subunit of the copolymerase required for synthesis of the backbone of HS, abolished binding to a greater extent than enzymatic removal of the chains with HSases (Fig. 4A and B), suggesting that the HSase treatment may underestimate the dependence on HS. Targeting *NDST1*, a GlcNAc *N*-deacetylase-*N*-sulfotransferase that N-deacetylates and N-sulfates *N*-acetylglucosamine residues, and *HS6ST1 and HS6ST2*, which introduces sulfate groups in the C6 position of glucosamine residues, significantly reduced binding (Figs. 4A and B). Although experiments with other sulfotransferases have not yet been done, the data suggests that S and RBD may recognize a specific pattern of sulfation.

**Figure 4.**
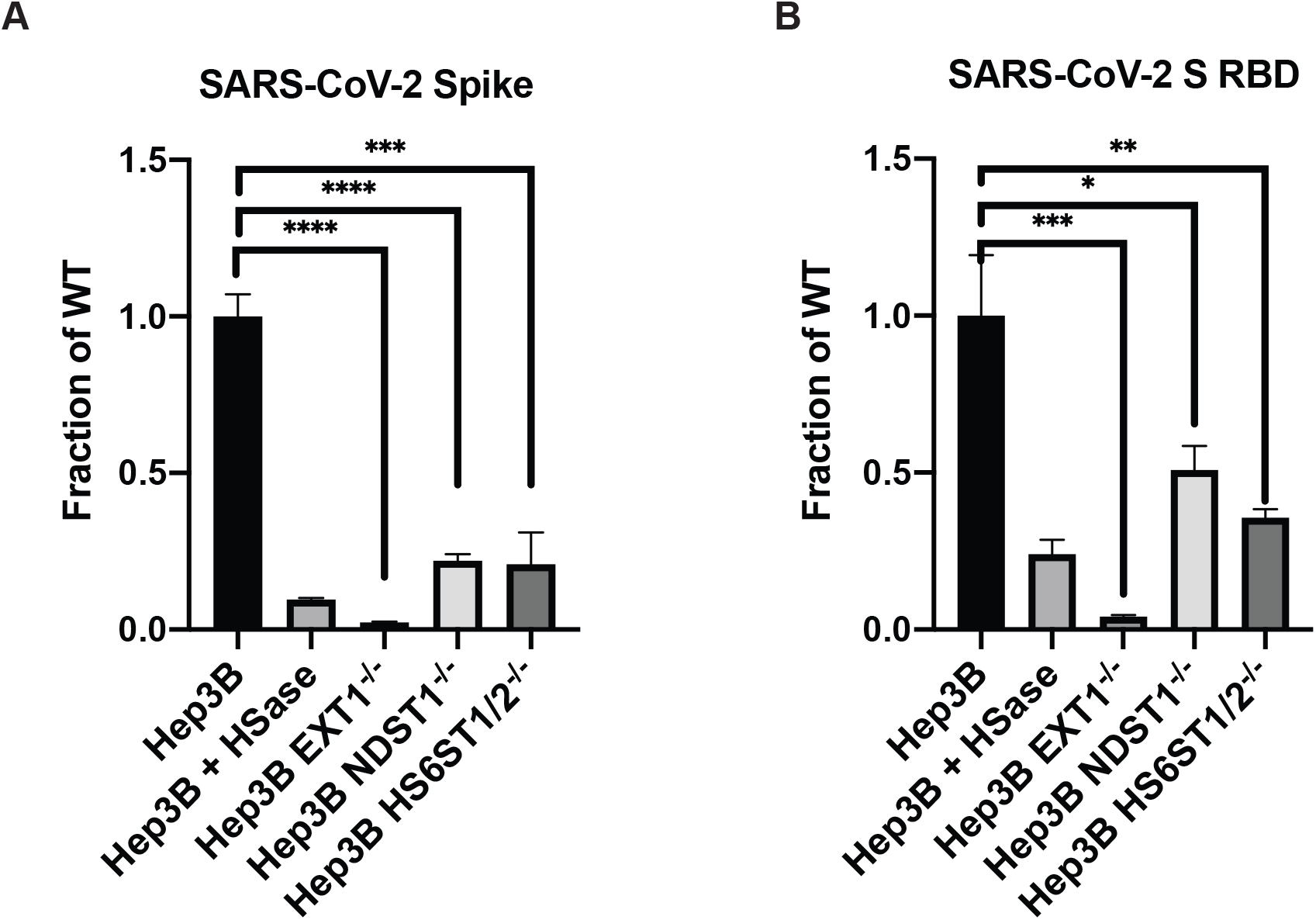
SARS-CoV-2 spike ectodomain protein binding to cellular heparan sulfate depends on sulfation. **A,** Binding of recombinant SARS-CoV-2 spike protein (20 μg/ml) to Hep3B mutants altered in HS biosynthesis enzymes. Specific enzymes that were mutated are listed along the x-axis. **B,** Binding of SARS-CoV-2 S RBD protein (20 μg/ml) Hep3B mutants. Binding was measured by flow cytometry. All experiments were repeated at least three times. Graphs shows representative experiments performed in technical triplicates. Statistical analysis by unpaired t-test. (ns: p > 0.05, *: p ≤ 0.05, **: p ≤ 0.01, ***: p ≤ 0.001, ****: p ≤ 0.0001).

### Heparin and heparan sulfates inhibit binding of spike ectodomain protein

To examine how variation in the HS structure affects binding, we isolated HS from human kidney, liver, lung and tonsil. The samples were depolymerized into disaccharides by treatment with HSases, and the disaccharides were then analyzed by LC-MS (Experimental Methods). The disaccharide analysis showed that lung HS has a larger proportion of *N*-deacetylated and *N*-sulfated glucosamine residues (grey bars) and more 2-*O*-sulfated uronic acids (green bars) than HS preparations from the other tissues (Fig. 5A). The different HS preparations also varied in their ability to block binding of RBD to H1299 cells (Fig 5B). Interestingly, HS isolated from lung was more potent compared to kidney and liver HS, consistent with the greater degree of sulfation of HS from this organ (Table 1). HS from tonsil was as potent as HS from lung, but the overall extent of sulfation was not as great, supporting the notion that the patterning of the sulfated domains in the chains may play a role in binding.

**Figure 5.**
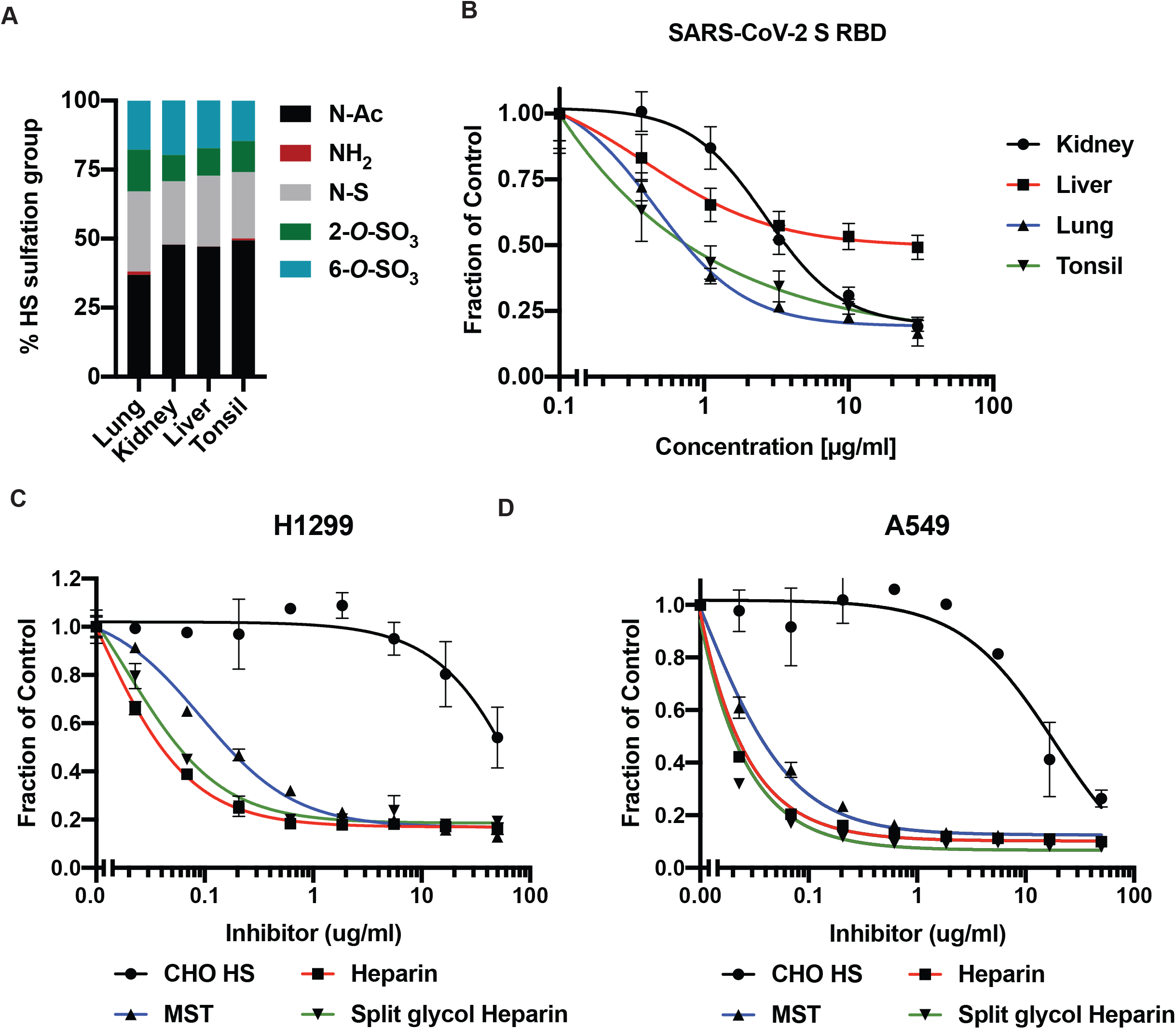
SARS-CoV-2 spike ectodomain protein binding to cells is differentially affected by HS from different organs and potently inhibited by heparinoids. **A,** LC-MS disaccharide analysis of HS isolated from human kidney, liver, tonsil, and lung tissue. **B,** Inhibition of binding of recombinant SARS-CoV-2 S RBD protein to H1299 cells, using tissue HS. Analysis by flow cytometry. **C,** inhibition of recombinant trimeric SARS-CoV-2 protein (20 μg/ml) binding to H1299 cells, using CHO HS, heparin, MST heparin, and split-glycol heparin. Analysis by flow cytometry. **D,** Similar analysis of A549 cells. Curve fitting was performed using non-linear fitting in Prism. IC_50_ values are listed in Table 1. Graphs shows representative experiments performed in technical duplicates or triplicates. (ns: p > 0.05, *: p ≤ 0.05, **: p ≤ 0.01, ***: p ≤ 0.001, ****: p ≤ 0.0001).

**Table 1.** IC_50_ values for heparin and HS as competitive inhibitors of S protein binding.

| **Cells** | **Inhibitor** | **IC<sub>50</sub> (µg/ml)** | **95% C.I. (µg/ml)** | **R<sup>2</sup> of fit** |
| --- | --- | --- | --- | --- |
| H1299 | CHO HS | 139 | 18 - ∞ | 0.803 |
|  | Heparin | 0.03 | 0.02 - 0.04 | 0.991 |
|  | MST Heparin | 0.12 | 0.09 - 0.15 | 0.991 |
|  | Split-Glycol Heparin | 0.04 | 0.03 - 0.06 | 0.971 |
|  | Kidney HS | 8.4 | 3.7 to 25 | 0.749 |
|  | Liver HS | 62 | 15 to ∞ | 0.627 |
|  | Lung HS | 2.1 | 0.78 to 5.8 | 0.828 |
|  | Tonsil HS | 2.5 | 0.74 to 7.5 | 0.838 |
| A549 | CHO HS | 19 | 8.6 - 49 | 0.907 |
|  | Heparin | 0.01 | 0.010 - 0.013 | 0.997 |
|  | MST Heparin | 0.03 | 0.026 - 0.032 | 0.997 |
|  | Split-Glycol Heparin | 0.01 | 0.007 to 0.008 | 0.999 |

Unfractionated heparin is derived from porcine mucosa and possesses potent anticoagulant activity due to the presence of a pentasaccharide sequence containing a crucial 3-*O*-sulfated *N*-sulfoglucosamine unit, which confers high affinity binding to antithrombin. Heparin is also very highly sulfated compared to HS with an average negative charge of –3.4 per disaccharide (the overall negative charge density of typical HS is –1.8 to –2.2 per disaccharide). MST cells, which were derived from a murine mastocytoma, make heparin-like HS that lacks the key 3-*O*-sulfate group and anticoagulant activity (Gasimli et al., 2014; Montgomery et al., 1992). The anticoagulant properties of heparin can also be removed by periodate oxidation, which oxidizes the vicinal hydroxyl groups in the uronic acids, resulting in what is called split-glycol heparin (Casu et al., 2004). All of these agents significantly inhibited binding of the S protein to H1299 and A549 cells (Fig 5C and 5D) yielding IC_50_ values in the range of 0.01-0.12 μg/ml (Table 1). Interestingly, the lack of 3-*O*-sulfation, crucial for the anticoagulant activity of heparin, had little effect on its inhibition of S binding. In contrast, CHO cell HS (containing 0.8 sulfates per disaccharide) only weakly inhibited binding (IC50 values of 18 and 139 μg/ml for A549 and H1299, respectively) (Table 1). These data suggest that inhibition by heparinoids is most likely charge dependent and independent of anticoagulant activity *per se*.

### Binding of S protein to ACE2 also depends on heparan sulfate

The experiments shown in Figs. 2C and 2D indicate that the binding sites in the RBD for ACE2 and HS function independently when tested as purified components. To explore if they function independently in cells, we genetically varied the expression of ACE2. Several attempts were made to measure ACE2 levels by Western blotting or flow cytometry with different mAbs and polyclonal antibodies, but a reliable signal was not obtained in any of the cell lines. However, expression was observed by RT-qPCR (Suppl. Fig. 3). However, when A375 cells were transfected with ACE2 cDNA, robust expression was detected by Western blotting (Fig. 6A). Binding of S ectodomain protein increased ~4-fold in the transfected cells (Fig. 6B), but the enhanced binding remained HS-dependent, as illustrated by the loss of binding after HSase-treatment. A375 cells carrying a CRISPR/Cas9 mediated deletion in the *B4GALT7* gene, which is required for glycosaminoglycan assembly (Fig. 6A, Suppl. Fig. S4), showed a similar dependence of spike binding on HS despite the overexpression of ACE2 in *B4GALT7*^-/-^ cells (Fig. 6B). To explore the impact of diminished ACE2 expression, we examined spike protein binding to A549 cells and in two CRISPR/Cas9 gene targeted clones C3 and C6 bearing biallelic mutations in ACE2 (Suppl. Fig. S4). Binding of S ectodomain protein was greatly reduced in the *ACE2*^-/-^ clones and the residual binding was sensitive to HSases (Fig. 6C). These findings show that binding of spike protein on cells involves cooperation between ACE2 and HS receptors, in contrast to their independent binding observed with purified components (Fig. 2).

**Figure 6.**
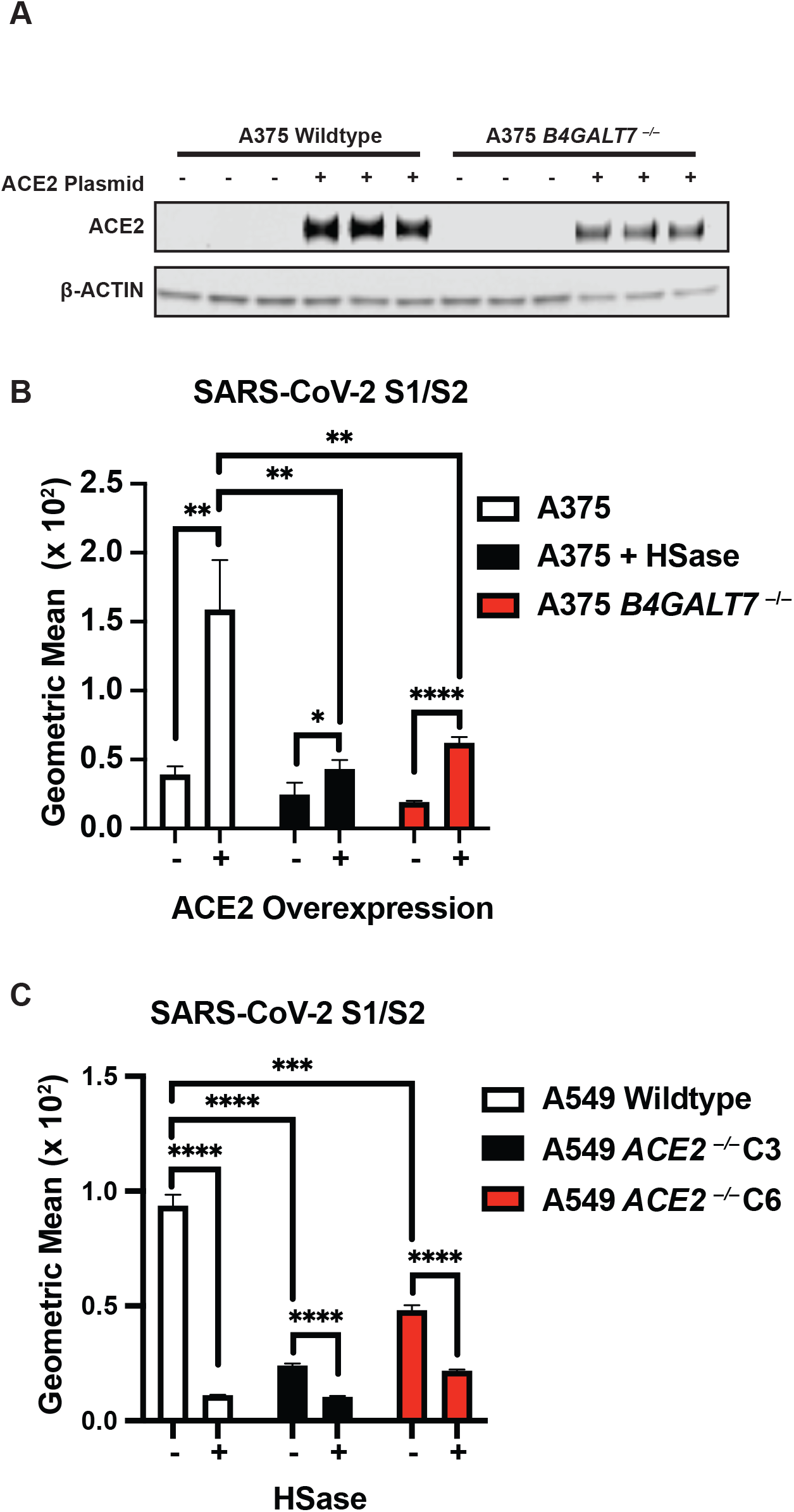
ACE2 and cellular heparan sulfate are both necessary for binding of SARS-CoV-2 spike ectodomain. **A,** Western blot shows overexpression of ACE2 in the A375 and A375 *B4GALT7*^-/-^ cells. A representative blot is shown. **B,** Binding of SARS-CoV-2 spike protein to cells with and without ACE2 overexpression. Note that binding is reduced in the mutants deficient in HS. **C,** Gene targeting of ACE2 in A549 using CRISPR/Cas9. The bars show spike binding to two independent ACE2 CRISPR/Cas9 knockout clones with and without HSase treatment. Note that binding depends on ACE2 expression and that residual binding depends in part on HS. All analyses were done by flow cytometry. The graphs show representative experiments performed in triplicate technical replicates. Statistical analysis by unpaired t-test. (ns: p > 0.05, *: p ≤ 0.05, **: p ≤ 0.01, ***: p ≤ 0.001, ****: p ≤ 0.0001).

### VSV S-protein pseudotyped virus infection depends on heparan sulfate

Assays using recombinant proteins do not recapitulate the multivalent presentation of the S protein as it occurs on the virion membrane. Thus, to extend these studies, pseudotyped vesicular stomatitis virus (VSV) was engineered to express the full-length SARS-CoV-2 S protein and GFP or luciferase. Vero E6 cells are commonly used in the study of SARS-CoV-2 infection, due to their high susceptibility to infection. Spike protein binding to Vero cells also depends on cellular HS, and binding was sensitive to HSases, heparin and split-glycol heparin (Fig. 7A). Interestingly, HSase treatment reduced binding to a lesser extent than the level of reduction observed in A549, H1299 and Hep3B cells. Vero cells express very high levels of ACE2 in comparison to other cells by Western blotting (Fig 7B) and by RT-qPCR (Suppl. Fig. S3), which may explain the decreased sensitivity to HSase. Infection of Vero cells by GFP-expressing VSV S protein pseudotyped virus was readily assessed qualitatively by fluorescence microscopy (inset) and quantitatively by flow cytometry (Fig. 7C and 7D). HSase treatment reduced infection ~3-fold. Infection by luciferin-expressing VSV S protein pseudotyped virus provided greater sensitivity, allowing studies with both high and low infection rates (Fig 7E, and 7F). Under either condition, infection was reduced 2 to 3-fold by HSase. Heparin very potently reduced infection more than ~4-fold at 0.5 μg/mL and higher concentrations (Fig. 7G). In contrast, studies of SARS-CoV-1 S protein pseudotype virus showed that HSase-treatment actually increased SARS-CoV-1 infection by more than 2-fold, suggesting that HS might interfere with binding of SARS-CoV-1 to ACE2 (Fig 7H). Infection of H1299 and A549 cells by SARS-CoV-2 S pseudotype virus was too low to obtain accurate measurements, but infection of Hep3B cells could be readily measured (Fig. 7I). HSase and mutations in *EXT1* and *NDST1* dramatically reduced infection 6- to 7-fold. Inactivation of the 6-*O*-sulfotransferases had at best a mild effect unlike its diminutive effect on S protein binding (Fig. 4), possibly due to the high valency conferred by multiple copies of S protein on the pseudovirus envelope.

**Figure 7:**
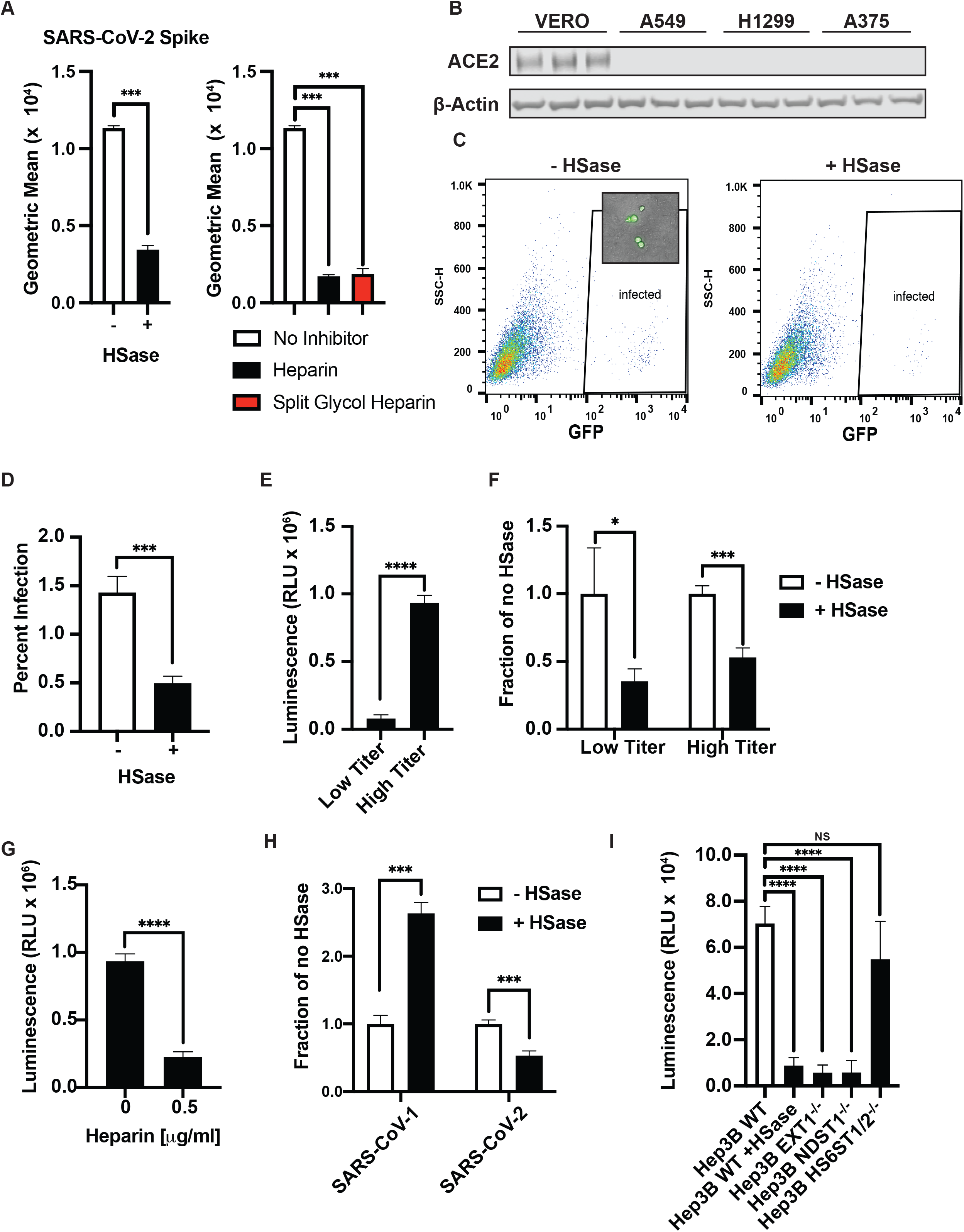
SARS-CoV-2 psuedovirus infection depends on heparan sulfate. **A,** Left panel, SARS-CoV-2 spike protein (20 μg/ml) binding to Vero cells measured by flow cytometry with and without HSase. Right panel, heparin and split-glycol heparin inhibit SARS-CoV-2 spike protein (20 μg/ml) binding to Vero cells by flow cytometry. Statistical analysis by unpaired t-test. **B**, Western Blot analysis of ACE2 expression in Vero E6 cells compared to A549, H1299 and Hep3B. A representative blot of three extracts is shown for each strain. **C,** Infection of Vero E6 cells with SARS-CoV-2 spike protein expressing pseudotyped virus expressing GFP. Infection was done with and without HSase treatment of the cells. Insert shows GFP expression in the infected cells by imaging. Counting was performed by flow cytometry with gating for GFP positive cells as shown. **D,** Quantitative analysis of GFP positive cells. **E,** Infection of Vero E6 cells with SARS-CoV-2 spike protein pseudotyped virus expressing luciferin. Infection was tittered and infection was measured by the addition of Bright-Glo^™^ and detection of luminescence. The figure shows infection experiments done at low and high titer. **F,** HSase treatment diminishes infection by SARS-CoV-2 spike protein pseudotyped virus (luciferase) at low and high titer. **G,** Heparin (0.5 μg/ml) blocks infection with SARS-CoV-2 spike protein pseudotyped virus (luciferase). **H,** Effect of HSase treatment of Vero E6 cells on the infection of both SARS-CoV-1 S and SARS-CoV-2 spike protein pseudotyped virus expressing luciferin. **I,** Infection of Hep3B with and without HSase and in Hep3B cells containing mutations in *EXT1, NDST1*, and *HS6ST1/HS6ST2*. Cells were infected with SARS-CoV-2 spike protein pseudotyped virus expressing luciferase. All experiments were repeated at least three times. Graphs shows representative experiments performed in technical triplicates. Statistical analysis by unpaired t-test. (ns: p > 0.05, *: p ≤ 0.05, **: p ≤ 0.01, ***: p ≤ 0.001, ****: p ≤ 0.0001).

### Cellular heparan sulfate is required for efficient infection by native SARS-CoV-2 virus

These studies were then extended to SARS-CoV-2 virus infection using strain USA-WA1/2020. Infection of Vero E6 cells was monitored by double staining of the cells with antibodies against the SARS-CoV-2 nucleocapsid and S proteins (Fig. 8A). Cells were treated with virus for 1 hr and the extent of infection was assayed 24 hrs later. Titration of the virus yielded infection rates of about 15-50%. Under either condition, treatment of the cells with HSases caused a ~5-fold reduction in the percentage of infected cells. Composite data from five separate experiments done in triplicate are shown in Fig. 8B (color coded for individual experiments). Data normalized to the values obtained in the absence of any treatment (mock) is shown in Fig. 8C. Titration of unfractionated heparin showed dose-dependent inhibition of infection (Fig. 8B and 8C) and emphasizes the potential for using unfractionated heparin or other non-anticoagulant heparinoids to prevent viral attachment.

**Figure 8.**
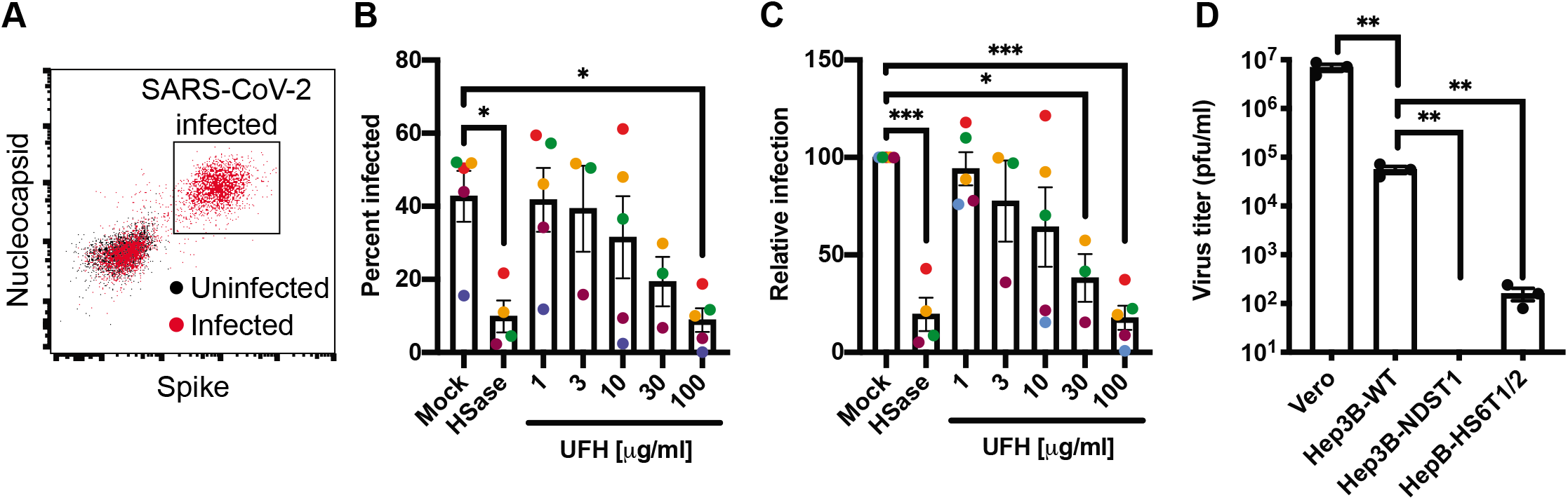
Manipulation of cellular heparan sulfate decreases infection of native SARS-CoV-2 virus. **A**, Flow cytometry analysis of SARS-CoV-2 infected (Red) or uninfected (Black) Vero cells stained with antibodies against SARS-CoV-2 nucleocapsid and spike protein. **B,** SARS-CoV-2 infection of Vero cells performed in the absence and presence of HSase, or with incubation with different concentrations of UFH. The extent of infection was analyzed by flow cytometry as in panel A. The graph shows a composite of five separate experiments performed in triplicate. The mean data from the individual experiments are colorized to allow for separate visualization **C,** Same data as in **B**, but with each experimental repeat normalized to the Mock infection. **D**, SARS-CoV-2 infection of Hep3B mutants altered in HS biosynthesis enzymes. Cells were infected for 1hr and incubated 48hrs, allowing for new virus to be formed. The resulting titers were determined by plaque assays on Vero E6 cells. Average values with standard error mean are shown, along with the individual data points. Statistical analysis by one-way ANOVA (**B, C**) or unpaired t-test (**D**); ns: *p* > 0.05, *: *p* ≤ 0.05, **: *p* ≤ 0.01, ***: *p* ≤ 0.001, ****: *p* ≤ 0.0001)

These findings were then extended to Hep3B cells and mutants altered in HS biosynthesis using a viral plaque assay. Virus was added to wildtype, *NDST1*^-/-^ and *HS6ST*1/2^-/-^ cells for 2 hr, the virus was removed, and after 2 days incubation a serial dilution of the conditioned culture medium was added to monolayers of Vero E6 cells. The number of plaques were then quantitated by staining and visualization. As a control, culture medium from infected Vero E6 cells was tested, which showed robust viral titers. Hep3B cells also supported viral replication, but to a lesser extent than Vero cells. Inactivation of *NDST1* in Hep3B cells abolished virus production, whereas inactivation of *HS6ST1/2*^-/-^ reduced infection about 3-fold (Fig. 8D). These findings demonstrate the requirement of cellular HS in mediating SARS-CoV-2 infection and replication.

## Discussion

In this report, we provide compelling evidence that HS is a necessary host attachment factor that mediates SARS-CoV-2 infection of various target cells. The receptor binding domain of the SARS-CoV-2 S protein binds to heparin, most likely through a docking site composed of positively charged amino acid residues aligned in a subdomain of the RBD that is separate from the site involved in ACE2 binding (Fig. 1A). Competition studies, enzymatic removal of HS, and genetic studies confirm that the S protein, whether presented as a recombinant protein (Figs. 2–6), in a pseudovirus (Fig. 7), or in intact SARS-CoV-2 virions (Fig. 8), binds to cell surface HS in a cooperative manner with ACE2 receptors. This data provides crucial insights into the pathogenic mechanism of SARS-CoV-2 infection and suggests HS-spike protein complexes as a novel therapeutic target to prevent infection.

The glycocalyx is the first point of contact for all pathogens that infect animal cells, and thus it is not surprising that many viruses exploit glycans, such as HS, as attachment factors. For example, the initial interaction of herpes simplex virus with cells involves binding to HS chains on one or more HS proteoglycans (Shieh et al., 1992; WuDunn and Spear, 1989) through the interactions with the viral glycoproteins gB and gC. Viral entry requires the interaction of a specific structure in HS with a third viral glycoprotein, gD (Shukla et al., 1999), working in concert with membrane proteins related to TNF/NGF receptors (Montgomery et al., 1996). Similarly, the human immunodeficiency virus binds to HS by way of the V3 loop of the viral glycoprotein gp120 (Roderiquez et al., 1995), but infection requires the chemokine receptor CCR5 (Deng et al., 1996; Dragic et al., 1996). Other coronaviruses also utilize HS, for example NL63 (HCoV-NL63) binds HS via the viral S protein in addition to ACE2 (Lang et al., 2011; Milewska et al., 2018; Milewska et al., 2014; Naskalska et al., 2019). In these examples, initial tethering of virions to the host cell plasma membrane appears to be mediated by HS, but infection requires transfer to a proteinaceous receptor. The data presented here shows that SARS-CoV-2 requires HS in addition to ACE2. Binding to heparin and ACE2 occurs independently in biochemical assays with purified components, but on the surface of cells the interaction apparently occurs in a co-dependent manner. We imagine a model in which cell surface HS acts as a “collector” to allow for optimal presentation to ACE2, making viral infection more efficient. HS varies in structure across cell types and tissues, as well as with gender and age (de Agostini et al., 2008; Feyzi et al., 1998; Ledin et al., 2004; Vongchan et al., 2005; Warda et al., 2006; Wei et al., 2011). Thus, it is possible HS contributes to the tissue tropism and the susceptibility of different patient populations in addition to levels of expression of ACE2 (Li et al., 2020).

Coronaviruses can utilize a diverse set of glycoconjugates as attachment factors. Human coronavirus OC43 (HCoV-OC43) and bovine coronavirus (BCoV) bind to 5-*N*-acetyl-9-*O*-acetylneuraminic acid (Hulswit et al., 2019; Tortorici et al., 2019), middle east respiratory syndrome virus (MERS) binds 5-*N*-acetyl-neuraminic acid (Park et al., 2019), and guinea fowl coronavirus binds biantennary di-*N*-acetyllactosamine or sialic acid capped glycans (Bouwman et al., 2019). Whether SARS-CoV-2 S protein binds to sialic acid remains unclear. Mapping the binding site for sialic acids in other coronavirus S proteins has proved elusive, but modeling studies suggest a location distinct from the HS binding site shown in Fig. 1 (Park et al., 2019; Tortorici et al., 2019). The S protein in murine coronavirus contains both a hemagglutinin domain for binding and an esterase domain that cleaves sialic acids that aids in the liberation of bound virions (Rinninger et al., 2006; Smits et al., 2005). Whether SARS-CoV-2 S protein, another viral envelope protein, or a host protein contributes an HS-degrading activity to aid in the release of newly made virions is unknown.

The repertoire of proteins in organisms that bind to HS make up the so called “HS interactome” and consists of a variety of different HS-binding proteins (HSBPs) (Xu and Esko, 2014). Unlike lectins that have a common fold that helps define the glycan binding site, HSBPs do not exhibit a conserved motif that allows accurate predictions of binding sites based on primary sequence. Instead, the capacity to bind heparin appears to have emerged through convergent evolution by juxtaposition of several positively charged amino acid residues arranged to accommodate the negatively charged sulfate and carboxyl groups present in the polysaccharide, and hydrophobic and H-bonding interactions stabilize the association. The RBD domains from the SARS-CoV-1 and SARS-CoV-2 S proteins are highly similar in structure (Fig. 1G), but the electropositive surface in SARS-CoV-1 S RBD is not as pronounced in SARS-CoV-2 S RBD (Figure 1H). *A priori* we predicted that the evolution of the HS binding site in the SARS-CoV-2 S protein might have occurred by the addition of arginine and lysine residues to its ancestor, SARS-CoV-1. Instead, we observed that 4 of the six predicted positively charged residues that make up the heparin-binding site are present in SARS-CoV-1 as well as most of the other amino acid residues predicted to interact with heparin (Fig. 1). SARS-CoV-1 has been shown to interact with cellular HS in addition to its entry receptors ACE2 and transmembrane protease, serine 2 (TMPRSS2) (Lang et al., 2011). Our analysis suggests that the putative heparin-binding site in SARS-CoV-2 S may mediate an enhanced interaction with heparin compared to SARS-CoV-1, and that this change evolved through as few as two amino acid substitutions, Thr444Lys and Glu354Asn.

It has previously been reported that SARS-CoV-2 infects target cells by interacting with ACE2 and that a possible reason for its higher virulence compared to SARS-CoV-1 may owe to an increase in ACE2 binding affinity (Hoffmann et al., 2020). Here, we showed that infection of SARS-CoV-1 S protein pseudotyped virus is actually increased upon heparin lyase-treatment (Fig. 7H). Thus, it is possible that the increased virulence of SARS-CoV-2 can in part be attributed to acquiring the capacity for more robust adhesion to HS.

The ability of heparin and HS to compete for binding of the SARS-CoV-2 S protein to cell surface HS and the inhibitory activity of heparin towards viral uptake of pseudovirus and native SARS-CoV-2 illustrates the therapeutic potential of agents that target the virus-HS interaction to control infection and transmission of SARS-CoV-2. There is precedent for targeting protein-glycan interactions as therapeutic agents. For example, Tamiflu targets influenza neuraminidase, thus reducing viral transmission, and sialylated human milk oligosaccharides can block sialic acid-dependent rotavirus attachment and subsequent infection in infants (Hester et al., 2013; von Itzstein, 2007). COVID-19 patients typically suffer from thrombotic complications ranging from vascular micro-thromboses, venous thromboembolic disease and stroke and often receive unfractionated heparin or low molecular weight heparin (Thachil, 2020). The findings presented here and in recent preprints uploaded to BioRxiv, suggests that both of these agents can block viral infection (Courtney Mycroft-West, 2020; Kim et al., 2020; Liu et al., 2020; Mycroft-West et al., 2020; Tandon et al., 2020). Thus, heparin may have dual action: as a hemostatic agent to prevent clotting and as an antiviral to decrease viral attachment. The anticoagulant activity of heparin, which is typically absent in HS, is not critical for its antiviral activity based on the observation that MST derived heparin and split-glycol heparin is nearly as potent as therapeutic heparin (Figs. 5 and 7). Additional studies are needed to address the potential overlap in the dose response profiles for heparin as an anticoagulant and antiviral agent. Antibodies directed to heparan sulfate or the binding site in the RBD might also prove useful for attenuating infection.

In conclusion, this work revealed HS as a novel receptor for SARS-CoV-2 and suggests the possibility of using HS mimetics, HS degrading lyases, and metabolic inhibitors of HS biosynthesis for the development of therapy to combat COVID-19.

## Methods

### Reagents

SARS-CoV-2 (2019-nCoV) spike Protein (RBD, His Tag) (Sino Biologicals, 40592-V08B), SARS-CoV-2 (2019-nCoV) spike S1 + S2 Protein (ECD, His Tag) (Sino Biologicals, 40589-V08B1), and SARS-CoV-2 spike protein (ECD, His & Flag Tag) (GenScript Z03481). Proteins were biotinylated using EZ-Link™ Sulfo-NHS-Biotin, No-Weigh™ (Thermo Fisher Scientific, A39256). Tag cleavage reagents were Pierce™ Tag Cleavage Enzymes, HRV 3C Protease Solution Kit (Thermo Scientific, 88946). Heparin lyases were from IBEX pharmaceuticals, Heparinase I (IBEX, 60-012), Heparinase II (IBEX, 60-018), Heparinase III (IBEX, 60-020). Protein production reagents included, Pierce™ Protein Concentrators PES Thermo Scientific™, Pierce™ Protein Concentrator PES (Thermo Fisher Scientific, 88517) and Zeba™ Spin Desalting Columns, 40K MWCO, 0.5 mL (Thermo Fisher Scientific, 87766). Antibodies used were, anti-spike antibody [1A9] (GeneTex, GTX632604), anti-Nucleocapsid antibody (GeneTex, GTX135357), Anti-HS (Clone F58-10E4) (Fisher Scientific, NC1183789), and Anti-ACE2 (Cell signaling, 4355S). Secondary antibodies were, Anti-His-HRP (Genscript, A00612), Avidin-HRP (Biolegend, 405103), and Streptavadin-Cy5 (Thermo Fisher, SA1011). Luciferase activity was monitored by Bright-Glo^™^ (Promega, E2610). All cell culture medias and PBS where from Gibco. SPL heparin was used for binding studies. APP Heparin (Hikma Pharmaceuticals USA Inc.) and Enoxaparin (Winthrop U.S.) were used in competition experiments. Heparin hexadecasaccharides were obtained from Galen Laboratory Supplies (#HO16).

### Molecular Modeling

An electrostatic potential map of the SARS-CoV-2 spike protein RBD domain was generated from a crystal structure (PDB:6M17) and visualized using Pymol (version 2.0.6 by Schrödinger). A dp4 fully sulfated heparin fragment was docked to the SARS-CoV-2 spike protein RBD using the ClusPro protein docking server (https://cluspro.org/login.php) (Kozakov et al., 2013; Kozakov et al., 2017; Vajda et al., 2017). Heparin-protein contacts and energy contributions were evaluated using the Molecular operating environment (MOE) software (Chemical Computing Group).

### SARS-CoV-2 spike protein production

Recombinant SARS-CoV-2 spike protein, encoding residues 1-1138 (Wuhan-Hu-1; GenBank: MN908947.3) with proline substitutions at amino acids positions 986 and 987 and a “GSAS” substitution at the furin cleavage site (amino acids 682-682), was produced in ExpiCHO cells by transfection of 6 x10^6^ cells/ml at 37 °C with 0.8 μg/ml of plasmid DNA using the ExpiCHO expression system transfection kit in ExpiCHO Expression Medium (ThermoFisher). One day later the cells were refed, then incubated at 32 °C for 11 days. The conditioned medium was mixed with cOmplete EDTA-free Protease Inhibitor (Roche). Recombinant protein was purified by chromatography on a Ni^2+^ Sepharose 6 Fast Flow column (1 ml, GE LifeSciences). Samples were loaded in ExpiCHO Expression Medium supplemented with 30 mM imidazole, washed in a 20 mM Tris-Cl buffer (pH 7.4) containing 30 mM imidazole and 0.5 M NaCl. Recombinant protein was eluted with buffer containing 0.5 M NaCl and 0.3 M imidazole. The protein was further purified by size exclusion chromatography (HiLoad 16/60 Superdex 200, prep grade. GE LifeSciences) in 20 mM HEPES buffer (pH 7.4) containing 0.2 M NaCl. Recombinant ectodomains migrate as a trimer assuming a monomer molecular mass of ~142,000 and 22 N-linked glycans per monomer (Watanabe et al., 2020). SDS-PAGE, TEM analysis, and SEC studies validate protein purity and the presence of trimers (Suppl. Fig. S1).

### SARS-CoV-2 spike RBD production

SARS-CoV-2 RBD (GenBank: MN975262.1; amino acid residues 319-529) was cloned into pVRC vector containing an HRV 3C protease-cleavable C-terminal streptavidin binding peptide (SBP)-His_8X_ tag and the sequence was confirmed. Recombinant protein was expressed by transient transfection of mammalian Expi293F suspension cells. Supernatants were harvested 5 days post-transfection and passed over Cobalt-TALON resin (Takara) followed by size exclusion chromatography on Superdex 200 Increase 10/300 GL (GE Healthcare) in PBS. Purity was assessed by SDS-PAGE analysis. Some initial optimization experiments utilized recombinant SARS-CoV-2 RBD and recombinant SARS-CoV-2 spike extracellular domain purchased from Sino Biological and Genscript. SDS-PAGE and SEC analysis is included in Suppl. Fig S1.

### Transmission electron microscopy

Samples of the recombinant trimeric spike protein ectodomain were diluted to 0.03 mg/ml in 1X TBS pH 7.4. Carbon coated copper mesh grids were glow discharged and 3 μL of the diluted sample was placed on a grid for 30 seconds then blotted off. Uniform stain was achieved by depositing 3 μL of uranyl formate (2%) on the grid for 55 seconds and then blotted off. Grids were transferred to a Thermo Fisher Morgagni operating at 80kV. Images at 56,000 magnification were acquired using a MegaView 2K camera via the RADIUS software. A dataset of 138 micrographs at 52,000x magnification and −1.5 μm defocus was collected on a FEI Tecnai Spirit (120keV) with a FEI Eagle 4k by 4k CCD camera. The pixel size was 2.06 Å per pixel and the dose was 25 e^-^/Å^2^. The Leginon (Suloway et al., 2005) software was used to automate the data collection and the raw micrographs were stored in the Appion (Lander et al., 2009) database. Particles on the micrographs were picked using DogPicker (Voss et al., 2009), stack with a box size of 200 pixels, and 2D classified with RELION 3.0 (Scheres, 2012).

### Recombinant ACE2 expression and purification

Secreted human ACE2 was transiently produced in suspension HEK293-6E cells. A plasmid encoding residues 1-615 of ACE2 with a C-terminal HRV-3C protease cleavage site, a TwinStrepTag and an His_8x_ Tag was a gift from Jason S. McLellan, University of Texas at Austin. Briefly, 100 mL of HEK293-6E cells were seeded at a cell density of 0.5 × 10^6^ cells/ml 24 hr before transfection with polyethyleneimine (PEI). For transfection, 100 μg of the ACE2 plasmid and 300 μg of PEI (1:3 ratio) were incubated for 15 min at room temperature. Transfected cells were cultured for 48 hr and fed with 100 mL fresh media for additional 48 hr before harvest. Secreted ACE2 were purified from culture medium by Ni-NTA affinity chromatography (Qiagen). Filtered media was mixed 3:1 (v/v) in 4X binding buffer (100 mM Tris-HCl, pH 8,0, 1,2 M NaCl) and loaded on to a self-packed column, pre-equilibrated with washing buffer (25 mM Tris-HCl, pH 8, 0.3 M NaCl, 20 mM imidazole). Bound protein was washed with buffer and eluted with 0.2 M imidazole in washing buffer. The protein containing fractions were identified by SDS-PAGE.

### Analytical Heparin-Sepharose Chromatography

SARS-CoV-2 spike protein in dPBS was applied to a 1-ml HiTrap heparin-Sepharose column (GE Healthcare). The column was washed with 5 ml of dPBS and bound protein was eluted with a gradient of NaCl from 150 mM to 1 M in dPBS.

### Biotinylation

For binding studies, recombinant spike protein and ACE2 was conjugated with EZ-Link^™^ Sulfo-NHS-Biotin (1:3 molar ratio; Thermo Fisher) in Dulbecco’s PBS at room temperature for 30 min. Glycine (0.1 M) was added to quench the reaction and the buffer was exchanged for PBS using a Zeba spin column (Thermo Fisher).

### Binding of spike protein to heparin

Heparin (APP Pharmaceuticals) (50 ng) in 5 μL solution each was pipetted into each well of a high binding plate. A set of wells were set up without heparin. A solution (0.2 mL) of 90% saturated (3.7 M) ammonium sulfate was added to each well and incubated overnight to immobilize the HS. The next day, the wells were washed twice with 0.2 mL of PBS then blocked with 0.2 mL of PBS containing 0.1% Tween 20 (PBST) and 0.1% BSA for 1 hr at room temperature. The wells were emptied and 45 μL of PBST/BSA with 0, 1, 3, 6, 10, 30, 60 or 100 nM of his/FLAG-tagged SARS-CoV-2 spike protein (GenScript, Z03481-100) was added to each well. The wells were incubated for 1 hr at room temperature, washed thrice with 0.2 mL of PBST, and then incubated with 45 μL each of 0.1 μg/mL of THE anti-his-HRP (GenScript, A00612) in PBST/BSA for 1 hr at room temperature. The wells were washed 5 times with 0.2 mL of PBST. TMB Turbo substrate was added to each well (0.1 mL), and the color was allowed to develop. The reaction was quenched by addition of 0.1 mL of 1 M sulfuric acid. The absorbance was measured by 450 nm.

### Immobilization and binding of ACE2, spike and heparin-BSA

High binding microtiter plates were coated with heparin-BSA (100 ng/well), ACE2 (150 ng/well), or S protein (200 ng/well) overnight at 4°C. The plates were then blocked for 3 hr at 37 °C with TSM buffer (20 mM Tris buffer, pH 7.4, containing 150 mM NaCl, 2 mM MgCl_2_, 2 mM CaCl_2_, 0.05% Tween-20, and 1% BSA) and a dilution series of biotinylated proteins prepared in TSM buffer was added to the plates in triplicate. Bound biotinylated protein was detected by adding Avidin-HRP (405103, BioLegend) diluted 1:2000 in TSM buffer. Lastly, the plates were developed with TMB turbo substrate for 5-15 min. The reaction was quenched using 1 M sulfuric acid and the absorbance was measured at 450 nm. To detect the formation of a ternary complex of ACE2, S protein and heparin-BSA, the plates were first coated with heparin BSA and incubated with S protein (100 nM). ACE2 binding was measured to bound spike protein as described above.

### Tissue Culture

NCI-H1299, A549, Hep3B, A375 and Vero E6 cells were from the American Type Culture Collection (ATCC). NCI-H1299 and A549 cells were grown in RPMI medium, whereas the other lines were grown in DMEM. Hep3B cells carrying mutations in HS biosynthetic enzymes were derived from the parent Hep3B line as described previously (Anower et al., 2019). All cells were supplemented with 10% (v/v) FBS, 100 IU/ml of penicillin and 100 μg/ml of streptomycin sulfate and grown under an atmosphere of 5% CO_2_ and 95% air. Cells were passaged at ~80% confluence and seeded as explained for the individual assays.

### Flow cytometry

Cells at 50-80% confluence were lifted with PBS containing 10 mM EDTA (Gibco) and washed in PBS containing 0.5% BSA. The cells were seeded into a 96-well plate at 10^5^ cells per well. In some experiments, a portion of the cells was treated with HSase mix (2.5 mU/ml HSase I, 2.5 mU/ml HSase II, and 5 mU/ml HSase III; IBEX) for 30 min at 37 °C in PBS containing 0.5% BSA. They were incubated for 30 min at 4°C with biotinylated spike protein (S1/S2 or RBD; 20 μg/ml or serial dilutions) in PBS containing 0.5% BSA. The cells were washed twice and then reacted for 30 min at 4°C with Streptavadin-Cy5 (Thermo Fisher; 1:1000 dilution) in PBS containing 0.5% BSA,. The cells were washed twice and then analyzed using a FACSCalibur or a FACSCanto instrument (BD Bioscience). All experiments were done a minimum of three separate times in three technical replicates. Data analysis was performed using FlowJo software and statistical analyses were done in Prism 8 (GraphPad).

### HS purification from tissues

Fresh human tissue was washed in PBS, frozen, and lyophilized. The dried tissue was crushed into a fine powder, weighed, resuspended in PBS containing 1 mg/mL Pronase (Streptomyces griseus, Sigma Aldrich) and 0.1% Triton X-100, and incubated at 37°C overnight with shaking. The samples were centrifuged at 20,000 x g for 20 min and the supernatant was mixed 1:10 with equilibration buffer (50 mM sodium acetate, 0.2 M NaCl, 0.1% Triton X-100, pH 6) and loaded onto a DEAE Sephacel column (GE healthcare) equilibrated with buffer. The column was washed with 50 mM sodium acetate buffer containing 0.2 M NaCl, pH 6.0, and bound GAGs were eluted with 50 mM sodium acetate buffer containing 2.0 M NaCl, pH 6.0. The eluate was mixed with ethanol saturated with sodium acetate (1:3, vol/vol) and kept at −20°C overnight, followed by centrifugation at 20,000 x g at 4°C for 20 min. The pellets were dried in a centrifugal evaporator and reconstituted in DNase buffer (50 mM Tris, 50 mM NaCl, 2.5 mM MgCl2, 0.5 mM CaCl2, pH 8.0) with 20 kU/mL bovine pancreatic deoxyribonuclease I (Sigma Aldrich) and incubated with shaking for 2 hr at 37°C. The samples were adjusted to 50 mM Tris and 50 mM NaCl, pH 8.0, and incubated for 4 hr at 37 °C with 20 mU/mL chondroitinase ABC (Proteus vulgaris, Sigma Aldrich). The HS was purified over a DEAE column and precipitated with 90% ethanol (Esko, 1993).

### HS digestion and MS analysis

For HS quantification and disaccharide analysis, purified HS was digested with a mixture of heparin lyases I-III (2 mU each) for 2 hr at 37 °C in 40 mM ammonium acetate buffer containing 3.3 mM calcium acetate, pH 7.0. The samples were dried in a centrifugal evaporator and reductively aminated with [^12^C_6_]aniline. The HS samples were analyzed by liquid chromatography/tandem mass spectrometry and quantified by inclusion of [^13^C6]aniline-tagged standard HS disaccharides as described (Lawrence et al., 2008). The samples were separated on a reverse phase column (TARGA C18, 150 mm x 1.0 mm diameter, 5 μm beads; Higgins Analytical, Inc.) using 5 mM dibutylamine as an ion pairing agent (Sigma-Aldrich), and fragmentation ions were monitored in negative mode. Separation was performed using the same gradient, capillary temperature and spray voltage as described (Lawrence et al., 2008). The analysis was done on an LTQ Orbitrap Discovery electrospray ionization mass spectrometer (Thermo Scientific) equipped with an Ultimate 3000 quaternary HPLC pump (Dionex).

### ACE2 overexpression and immunoblotting

The ACE2 expression plasmid (Addgene, plasmid #1786) was received in bacteria and purified with a maxiprep kit (Zymogen). A375 wild-type and *B4GALT7*^-/-^ cells were transfected with 15 μg ACE2 expression plasmid in a mixture of DMEM, OptiMEM (Gibco), Lipofectamine 2000 with Plus reagent (Invitrogen). After 4 hr, the medium was replaced with DMEM/10% FBS and the cells were incubated for 3 d before being used for experiments.

To measure ACE2 expression, cells were lysed in RIPA Buffer (Millipore, 20-188) with protease inhibitors (cOmplete, Mini, EDTA-free Protease Inhibitor Cocktail, Roche, 11836170001). The lysates were incubated on ice for 1 hr and then centrifuged for 10 min at 10,000 x g at 4 °C. The supernate was collected and protein was quantified by BCA assay (Pierce BCA Protein Assay Kit ThermoFisher Scientific, 23225). To measure ACE2 expression in transfected cells, a protein ladder (PageRule Plus Prestained Protein Ladder, Thermo Scientific, PI26619) was mixed with 4 μg of each cell lysate in triplicate. To measure endogenous levels of hACE2 in cell lines, a protein ladder (Chameleon Duo Pre-stained Protein Ladder, Licor, 928-60000) was mixed with 15 μg of each cell lysate in triplicate. Samples were separated by electrophoresis on a 15-well Bolt 4-12% Bis-Tris gel (Invitrogen, NW04125 or Invitrogen, NP0336) in NuPAGE MOPS SDS Running Buffer (Invitrogen, NP0001). The gels were transferred onto a PVDF membrane (Immobilon-FL PVDF Membrane, Millipore, IPFL0010) in NuPAGE Transfer Buffer (Invitrogen, NP00061). The membranes were blocked 1 hr at room temperature with Odyssey PBS Blocking Buffer (Li-Cor, 927-40000) or in 5% Blotting-Grade Blocker (Biorad, 1706404) in TBST (50 mM Tris buffer, pH 7.5 containing 150 mM NaCl and 0.1% Tween-20) and then incubated overnight at 4 °C with anti-hACE2 antibody (1:1000; R&D AF933) and anti-beta actin (1:2000; CST 4970) in 5% BSA (Sigma-Aldrich A9647) in TBST. The blot was incubated at room temperature for 1 hr with appropriate secondary antibodies (Donkey anti-Goat, Li-Cor, 926-32214; IRDye 690LT Donkey anti-Rabbit, Li-Cor, 926-68023; both at 1:20,000) in 5% BSA or 2.5% Blotting-Grade Blocker and 0.02% SDS in TBST. The blots were imaged using an Odyssey Infrared Imaging System (Li-Cor) and quantified using the companion ImageStudio software.

### qPCR

mRNA was extracted from the cells using TRIzol (Invitrogen) and chloroform and purified using the RNeasy Kit (Qiagen). cDNA was synthesized from the mRNA using random primers and the SuperScript III First-Strand Synthesis System (Invitrogen). SYBR Green Master Mix (Applied Biosystems) was used for qPCR following the manufacturer’s instructions, and the expression of TBP was used to normalize the expression of *ACE2* between the samples. The qPCR primers used were as follows: *ACE2* (human) forward: 5’ – CGAAGCCGAAGACCTGTTCTA - 3’ and reverse: 5’ – GGGCAAGTGTGGACTGTTCC – 3’; and *TBP* (human) forward: 5’ – AACTTCGCTTCCGCTGGCCC – 3’ and reverse: 5’ – GAGGGGAGGCCAAGCCCTGA – 3’.

### Mutant cell line generation

To generate the Cas9 lentiviral expression plasmid, 2.5 x 10^6^ HEK293T cells were seeded to a 10-cm diameter plate in DMEM supplemented with 10% FBS. The following day, the cells were co-transfected with the psPAX2 packaging plasmid (Addgene, plasmid #12260), pMD2.g envelope plasmid (Addgene, plasmid #12259), and lenti-Cas9 plasmid (Addgene, plasmid #52962) in DMEM supplemented with Fugene6 (30μL in 600μL DMEM). Media containing the lentivirus was collected and used to infect A549 WT and A375 WT cells, which were subsequently cultured with 5 μg/mL and 2 μg/mL blasticidin, respectively, to select for stably transduced cells. A single guide RNA (sgRNA) targeting *ACE2* (5’-TGGATACATTTGGGCAAGTG −3’) and one targeting *B4GALT7* (5’-TGACCTGCTCCCTCTCAACG-3’) was cloned into the lentiGuide-Puro plasmid (Addgene plasmid #52963) following published procedure (Sanjana et al., 2014). The lentiviral sgRNA construct was generated in HEK293T cells, using the same protocol as for the Cas9 expression plasmid, and used to infect A549-Cas9 and A375-Cas9 cells to generate CRISPR knockout mutant cell lines. After infection, the cells were cultured with 2 μg/mL puromycin to select for cells with stably integrated lentivirus. After 7 d, the cells were serially diluted into 96-well plates. Single colonies where expanded and DNA was extracted using the DNeasy blood and tissue DNA isolation kit (Qiagen). Proper editing was verified by sequencing (Genewiz Inc.) and gene analysis using the online ICE tool from Synthego (Supplemental Fig 2).

### Preparation and infection by pseudotyped VSV

Vesicular Stomatitis Virus (VSV) pseudotyped with spike proteins of SARS-CoV-2 were generated according to a published protocol (Whitt, 2010). Briefly, HEK293T, transfected to express full length SARS-CoV-2 spike proteins, were inoculated with VSV-G pseudotyped ΔG-luciferase or GFP VSV (Kerafast, MA). After 2 hr at 37°C, the inoculum was removed and cells were refed with DMEM supplemented with 10% FBS, 50 U/mL penicillin, 50 μg/mL streptomycin, and VSV-G antibody (I1, mouse hybridoma supernatant from CRL-2700; ATCC). Pseudotyped particles were collected 20 hr post-inoculation, centrifuged at 1,320 × g to remove cell debris and stored at −80°C until use.

Cells were seeded at 10,000 cells per well in a 96-well plate. The cells (60-70% confluence) were treated with HSases for 30 min at 37 °C in serum-free DMEM. Culture supernatant containing pseudovirus (20-100 μL) was adjusted to a total volume of 100 μL with PBS, HSase mix or the indicated inhibitors and the solution was added to the cells. After 4 hr at 37 °C the media was changed to complete DMEM. The cells were then incubated for 16 hr to allow expression of reporter gene. Cells infected with GFP containing virus were visualized by fluorescence microscopy and counted by flow cytometry. Cells infected with Luciferase contaning virus were analyzed by Bright-Glo^™^ (Promega) using the manufacturers protocol. Briefly, 100 μL of luciferin lysis solution was added to the cells and incubated for 5 min at room temperature. The solution was transferred to a black 96-well plate and luminescence was detected using an EnSpire multimodal plate reader (Perkin Elmer). Data analysis and statistical analysis was performed in Prism 8.

### Infection by SARS-CoV-2 virus

SARS-CoV-2 isolate USA-WA1/2020 (BEI Resources, #NR-52281) was propagated and infectious units quantified by plaque assay using Vero E6 cells. The cells were treated with or without HSase mix (IBEX Pharmaceuticals) or with unfractionated heparin (UFH) and infected with SARS-CoV-2 for 1 hr at 37 °C. HSase-treated Vero E6 cells were incubated with HSase mix 30 min prior to infection until 24 hr post-infection or with UFH at the indicated concentrations from the start of infection until 24 hr post-infection. The cells were washed twice with PBS, lifted in Trypsin-EDTA (Gibco), and fixed in 4% formaldehyde for 30 min. Cells were permeabilized for flow cytometry using BD Cytofix/Cytoperm according to the manufacturers protocol for fixed cells and stained with anti-spike antibody [1A9] (GeneTex GTX632604) and anti-Nucleocapsid antibody (GeneTex GTX135357) that were directly conjugated with Alexa Fluor 647 and Alexa Fluor 594 labeling kits (Invitrogen), respectively. Cells were then analyzed using an MA900 Cell Sorter (Sony).

### Virus plaque assays

Confluent monolayers of Vero E6 or Hep3B cells were infected with SARS-CoV-2 at an MOI of 0.1. After one hour of incubation at 37 °C, the virus was removed, and the medium was replaced. After 48 hr, cell culture supernatants were collected and stored at −80°C. Virus titers were determined by plaque assays on Vero E6 monolayers. In short, serial dilutions of virus stocks in Minimum Essential Media MEM medium (Gibco, #41500-018) supplemented with 2% FBS was added to Vero E6 monolayers on 24-well plates (Greiner bio-one, #662160) and rocked for 1 hr at room temperature. The cells were subsequently overlaid with MEM containing 1% cellulose (Millipore Sigma, #435244), 2% FBS, and 10 mM HEPES buffer, pH 7.5 (Sigma #H0887) and the plates were incubated at 37 °C under an atmosphere of 5% CO2/95% air for 48 hr. The plaques were visualized by fixation of the cells with a mixture of 10% formaldehyde and 2% methanol (v/v in water) for 2 hr. The monolayer was washed once with PBS and stained with 0.1% Crystal Violet (Millipore Sigma # V5265) prepared in 20% ethanol. After 15 min, the wells were washed with PBS, and plaques were counted to determine the virus titers. All work with the SARS-CoV-2 was conducted in Biosafety Level-3 conditions either at the University of California San Diego or at the Eva J Pell Laboratory, The Pennsylvania State University, following the guidelines approved by the Institutional Biosafety Committees.

## Author Contributions

T.M.C., D.R.S., and J.D.E. conceived, initiated, and coordinated the project. T.M.C., D.R.S., C.B.S., C.D.P., J.P., B.E.T., J.T., C.A.G., A.N., S.A.M, G.J.G., and A.F.C. designed and performed the experimental work. Z.Y., X.Y., Y.P., J.F., B.H., T.C., S.K.C., R.P., K.G., A.W., A.G.S., and K.G. supplied reagents. B.P.K., C.M., J.J., K.D.C., S.L.L. and PL.S.M.G. provided essential discussion and advice. T.M.C., D.R.S., and J.D.E. performed the main fundraising. T.M.C., D.R.S., and J.D.E. wrote the manuscript. All authors discussed the experiments and results, read, and approved the manuscript.

## Conflicts of Interest

J.D.E. is a co-founder of TEGA Therapeutics. J.D.E. and The Regents of the University of California have licensed a University invention to and have an equity interest in TEGA Therapeutics. The terms of this arrangement have been reviewed and approved by the University of California, San Diego in accordance with its conflict of interest policies. C.A.G and B.E.T are employees of TEGA Therapeutics.

## Acknowledgements

We thank Eugene Yeo (UC San Diego), John Guatelli (UC San Diego), Mark Fuster (UC San Diego) and Stephen Schoenberger (La Jolla Institute for Immunology) for many helpful discussions, and Annamaria Naggi and Giangiacomo Torri from the Ronzoni Institute for generously providing split-glycol heparin. This work was supported by RAPID grant 2031989 from the National Science Foundation and Project 3 of NIH P01 HL131474 to J.D.E.; The Alfred Benzon Foundation to T.M.C; NIH R01 AI146779 and a Massachusetts Consortium on Pathogenesis Readiness MassCPR grant to A.G.S.; DOD grant W81XWH-20-1-0270 and Fluomics/NOSI U19 AI135972 to S.K.C; a Career Award for Medical Scientists from the Burroughs Wellcome Fund to A.F.C.; Bill and Melinda Gates Foundation grant OPP1170236 to A.B.W.; COVID19 seed funding from the Huck Institutes of the Life Sciences and Penn State start-up funds to J.J.; and T32 training grants GM007753 for B.M.H. and T.C and AI007245 for J.F.

## Supplemental Figures

**Supplemental Figure S1. Location of the putative heparin/HS binding site in the spike protein RBD from SARS-CoV-2.** PDB files 6VSB and 6M0J were used to model the spike protein. The residues colored pink on the three RBDs (444+509+346+354+356+357+355+466+ 347+348+349+353+450+448+451+352) make up a potential binding site for heparin and heparan sulfate.

**Supplemental Figure S2. A,** SDS-PAGE gel of recombinant SARS-CoV-2 spike ectodomain protein produced in ExpiCho cells and commercial recombinant SARS-CoV-2 RBD. **B,** Transmission electron micrographs of recombinant SARS-CoV-2 spike ectodomain protein. **C,** Size exclusion chromatography of recombinant SARS-CoV-2 spike ectodomain protein on a Superose 6 column. **D,** SDS-PAGE gel of recombinant SARS-CoV-2 RBD produced in ExpiHEK cells. **E,** Size exclusion chromatography of recombinant SARS-CoV-2 RBD on a Superdex200 column.

**Supplemental Figure S4. RT-qPCR analysis of ACE2 expression.**

**Supplemental Figure S4. DNA sequencing of ACE2 mutant alleles.**

