## Supplementary figures and images for "SARS-CoV-2 Infection Depends on Cellular Heparan Sulfate and ACE2"

### Supplemental Figure S1.

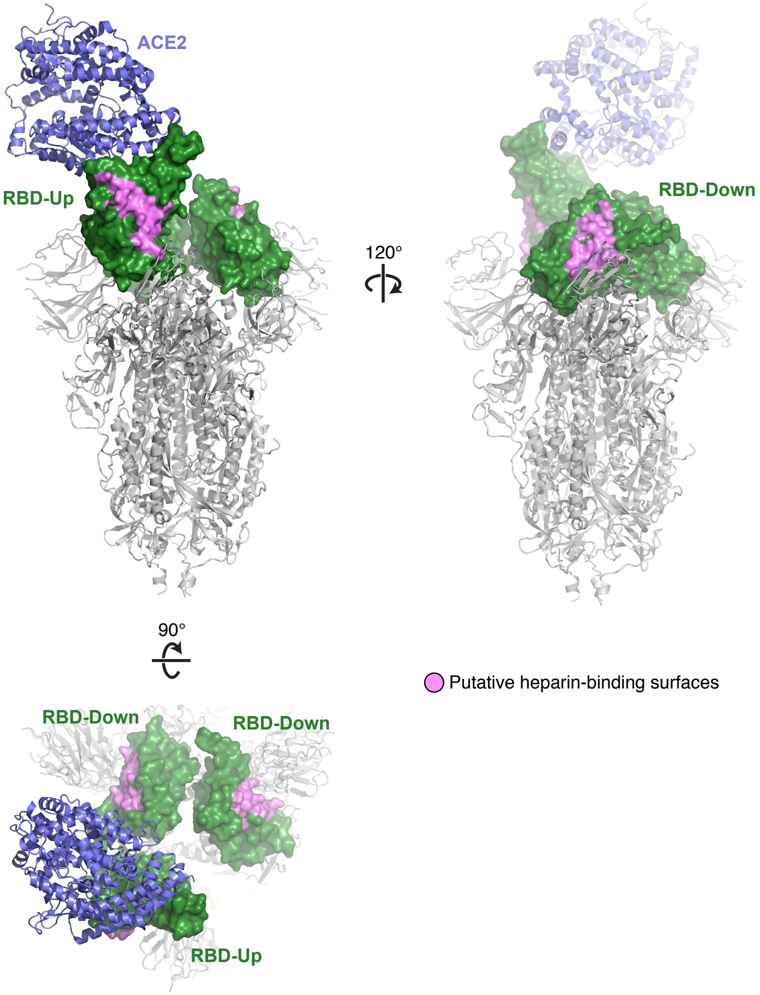

### Supplemental Figure S2.

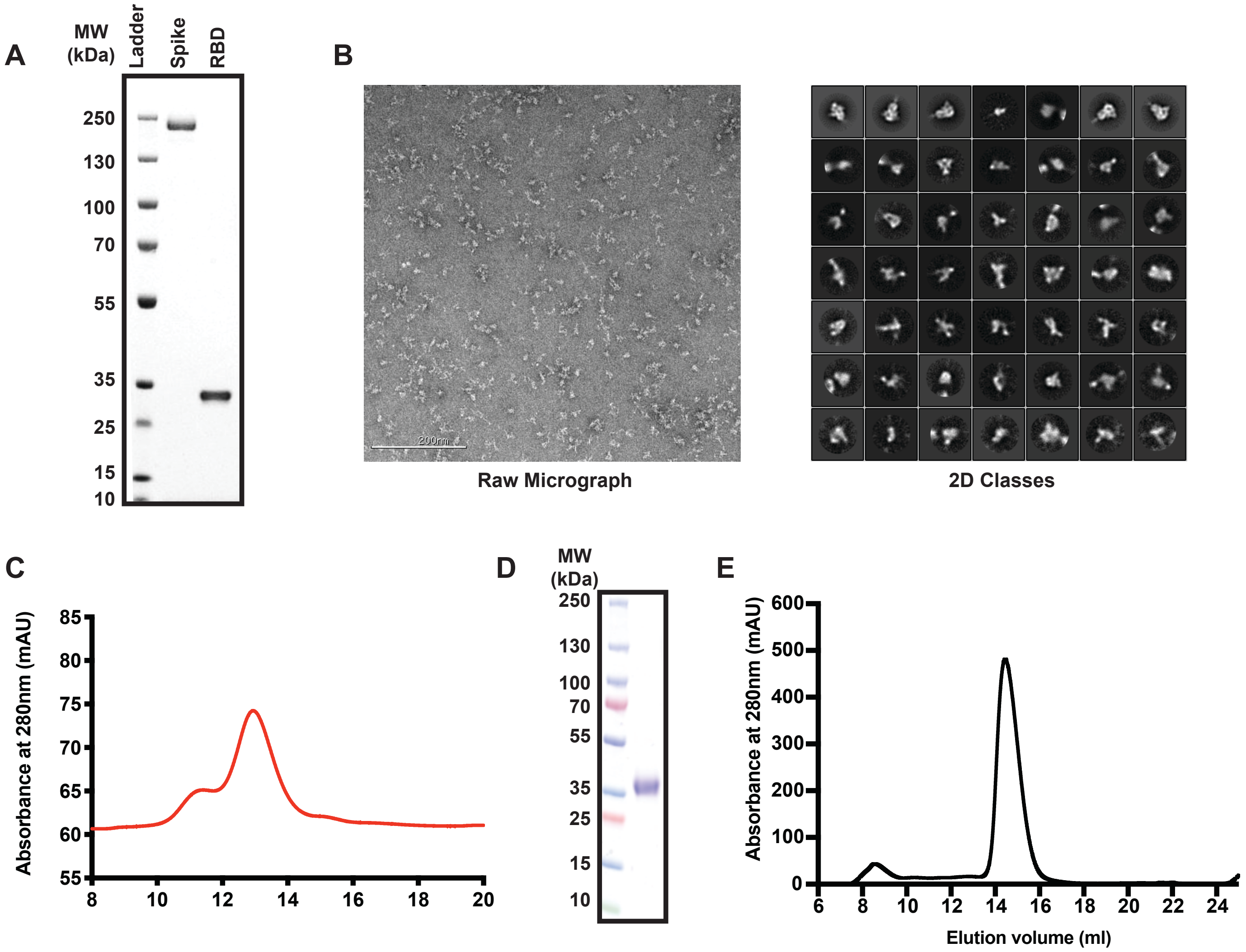

### Supplemental Figure S3.

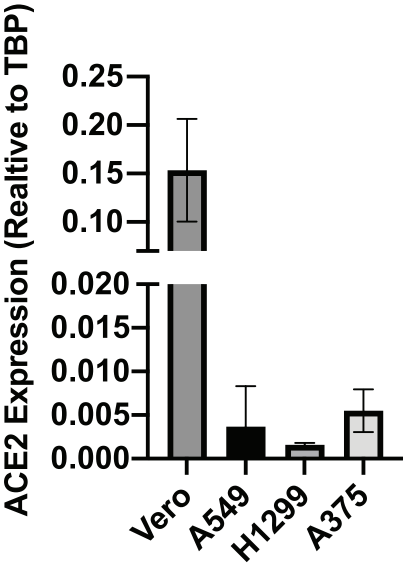

### Supplemental Figure S4.

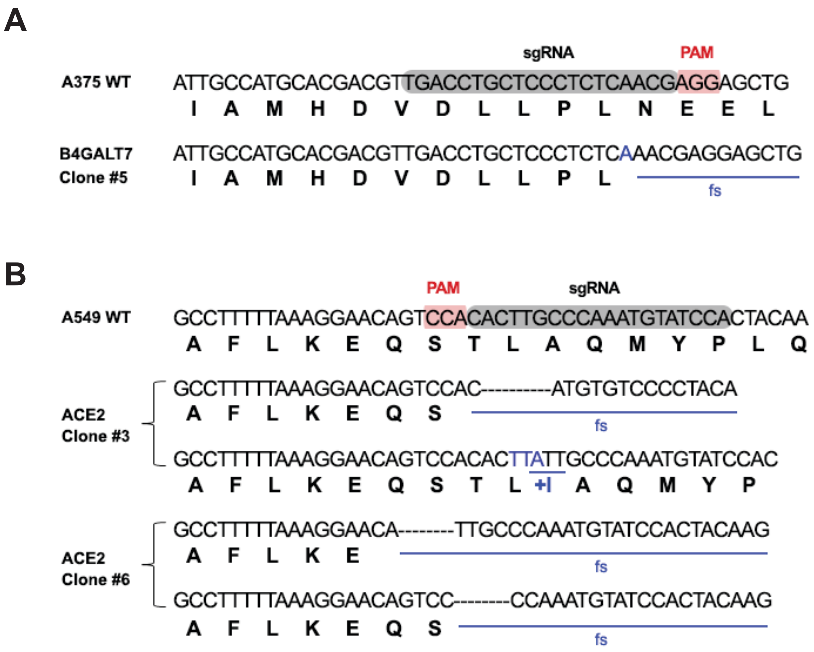
